# Length of uninterrupted CAG repeats, independent of polyglutamine size, results in increased somatic instability and hastened age of onset in Huntington disease

**DOI:** 10.1101/533414

**Authors:** Galen E.B. Wright, Jennifer A. Collins, Chris Kay, Cassandra McDonald, Egor Dolzhenko, Qingwen Xia, Kristina Bečanović, Alicia Semaka, Charlotte M. Nguyen, Brett Trost, Fiona Richards, Emilia K. Bijlsma, Ferdinando Squitieri, Stephen W. Scherer, Michael A. Eberle, Ryan K.C. Yuen, Michael R. Hayden

## Abstract

Huntington disease (HD) is an autosomal dominant neurological disorder that is caused by a CAG repeat expansion, translated into polyglutamine, in the huntingtin (*HTT)* gene. Although the length of this repeat polymorphism is inversely correlated with age of onset (AOO), it does not fully explain the variability in AOO. Genomic studies have provided evidence for the involvement of DNA repair in modifying this trait, potentially through somatic repeat instability. We therefore assessed genetic variants within the 12bp interrupting sequence between the pathogenic CAG repeat and the adjacent polymorphic proline (CCG) tract in the *HTT* gene and identified variants that result in complete loss of interruption (LOI) between the *HTT* CAG and CCG repeats. Analysis of multiple HD pedigrees showed that this variant is associated with dramatically earlier AOO and is particularly relevant to HD patients with reduced penetrance alleles. On average AOO of HD is hastened by an average of 25 years in LOI carriers. This finding indicates that the number of uninterrupted CAG repeats is the most significant contributor to AOO of HD and is more impactful than polyglutamine length, which is not altered in these patients. We show that the LOI variant is associated with increases in both somatic and germline repeat instability, demonstrating a potential mechanism for this effect. Screening individuals from the general population (*n*=2,674 alleles) suggests that the variant occurs only in expanded CAG repeat alleles. Identification of this modifier has important clinical implications for disease management of HD families, especially for those with genotypes in the reduced penetrance ranges.

## INTRODUCTION

The age of clinical onset (AOO) of Huntington disease (HD) in the past has been chiefly determined by the length of the expanded CAG repeat, translated into polyglutamine, in the *HTT* gene.^1^ However, this repeat polymorphism does not fully explain the variability in AOO, and HD patients with identical expanded repeat lengths frequently present with clinical symptoms at different ages.^2^ Predictive testing in HD is particularly inadequate for carriers of reduced penetrance (RP) alleles (36 - 39 CAGs), where the majority of carriers remain asymptomatic into old age and only a small proportion present with HD at some point in their lives.^3; 4^ Some HD patients with RP alleles manifest with much earlier ages and the reason underlying this variability at the same polyglutamine length remains unexplained. Furthermore, differences in AOO observed between HD patients have been shown to be influenced by heritable factors, suggesting that other genetic modifiers play an important role in modifying disease onset.^5; 6^ The ability to more accurately predict the AOO of HD is of great clinical value and has important implications for disease management in these individuals.

Recent studies have identified candidate modifier regions for HD onset, both at the *HTT* locus^7^ and across the genome.^8; 9^ Many of the novel HD modifier genes that have been identified are involved in DNA repair-related processes, potentially mediating the somatic instability of the pathogenic repeat.^8; 9^ Since the *HTT* repeat region is highly complex, and sequence variants in this area have previously been described,^5; 10-12^ we investigated whether genetic variants within the CAG-CCG region might influence AOO in HD patients and repeat instability.

In the current study, we demonstrate that sequence variants in the *HTT* gene, resulting from transitions (CAA to CAG) in the common interrupting sequence cause complete loss of interrupted CAGs in the pathogenic repeat, dramatically altering clinical onset in HD patients more than any other previously described modifier. The impact of this loss of interruption variant (LOI) is seen most obviously in subjects with CAG repeat lengths in the RP range and onset of HD is hastened by an average of 25 years in all manifest carriers. This configuration of variants leaves the polyglutamine and polyproline lengths unchanged at the protein level, but extends the uninterrupted CAG and CCG repeats. Further, in contrast to the effect of the LOI variant, we show that a distinct duplication of the common CAG repeat interruption delays HD onset. Semi-quantitative analysis of instability in human tissue shows that the LOI variant is associated with increased somatic and germline instability of the CAG repeat. These results suggest that altered somatic instability of the CAG repeat underlies the phenotypic effects of the LOI variant.

## METHODS

### Patient populations

Genomic DNA from HD patients was obtained from the HD Biobank at the University of British Columbia (UBC) or through collaborators from pedigrees of HD patients found to be carrying the LOI variant. All samples were collected, stored and accessed with informed consent and ethical approval from the UBC / Children’s and Women’s Health Centre of British Columbia Research Ethics Board (UBC C&W REB H06-70467 and H06-70410). AOO was determined by the clinicians treating the patients or ascertained from their medical records. The predicted AOO for HD patients based on CAG repeat length was calculated according to the Langbehn *et al.* formula,^13^ and AOO ratios (i.e., predicted / observed AOO), along with related percentiles were calculated as previously described.^7^

### HTT CAG and CCG repeat sizing and interrupting sequence characterization

CAG and CCG repeat sizing was performed with control samples of known repeat lengths, using previously described methods, at the Centre for Molecular Medicine and Therapeutics at UBC in Vancouver, Canada.^3; 14^ Haplotyping of single nucleotide polymorphisms spanning the *HTT* locus was also carried out as previously described.^15^ Variants in the interrupting sequence between the *HTT* CAG-CCG repeat tracts were genotyped by clonal sequencing. Briefly, polymerase chain reaction (PCR) products encompassing the *HTT* CAG-CCG repeat tracts (*HTT*-CAG-3-F-*Eco*RI: 5’-GATCGAATTCATTGCCCCGGTGCTGAGCG and *HTT*-CAG-3-R-*Hind*III: 5’-GATCAAGCTTGCGGGCCCAAACTCACGGTC) were cloned into pUC19 plasmids following restriction enzyme double digest (6 units of *Eco*R1 and *Hind*III) and ligation. Vectors were subsequently transformed into DH5-α *E. coli* cells and positive clones were identified via colony PCR, then cultured overnight for extraction with QIAprep Spin Miniprep Kits (Qiagen, Hilden, Germany) and Sanger sequencing with the M13-R primer (5’-CAGGAAACAGCTATGAC).

### Genotyping the HTT-LOI in general population controls

The frequency of the LOI variant was determined in a cohort of 1,657 unrelated general population controls, recruited as unaffected parents in an autism spectrum disorder study (a specific cohort within the Autism Speaks MSSNG Project).^16^ Sequence-graph based alignment of PCR-free whole-genome sequence data from these individuals was performed with ExpansionHunter (v3.0.0-rc1),^17^ which explicitly models the *HTT* CAG-CCG repeats and the interrupting sequence region. We restricted analyses to samples with CCG repeat calls within what has been detected from traditional fragment analysis sizing (i.e., CCG repeats between 5 and 12). For each interrupting sequence in each sample we calculated the ratio of observed reads that fully span the interruption to their expected per-haplotype number (O/E ratio).

### *HTT CAG somatic expansion ratio calculations* and *germline instability estimates*

Electropherogram traces from fluorescently-labelled CAG sizing PCR products were used to calculate an expansion index to measure the somatic instability of the pathogenic repeat, since similar approaches have been successfully employed to measure huntingtin CAG instability.^18^ Additional LOI carriers were identified by screening additional family members in the HD pedigrees described above. Reactions were performed in triplicate and PCR products were diluted (1:60) before being run on the ABI Prism 3130xl Genetic Analyzer using manufacturer protocols (Applied Biosystems, Foster City, CA). Traces were assigned using GeneMapper Software v4.0 (Thermo Fisher Scientific) and the expansion index was calculated using the area under all expanded CAG repeat lengths relative to the area under the most prominent peak. Small-pool PCR data for the *HTT* CAG repeat in sperm from 34 European ancestry subjects, were also analyzed to assess germline CAG instability, as described by Semaka *et al.*19.

### Statistical analyses

Statistical and bioinformatic analyses were performed in R. Significant differences between genotype groups with regards to AOO and related information were calculated using a Wilcoxon rank-sum test. Significant differences in log-transformed instability/expansion measures and LOI carrier status, CAG repeat length, and age were assessed using linear regression. Residuals were checked for normality with the Shapiro-Wilk test.

## RESULTS

### The HTT CAG-CCG LOI variant, resulting in an uninterrupted CAG tract, is associated with an earlier age of HD clinical onset

Analysis of the interrupting sequence identified a key HD modifier variant (Figure 1) that results in complete loss of the interrupted CAG repeats in the pathogenic repeat (i.e., CA**A**-CAG to CA**G**-CAG), without changing the length of the polyglutamine tract. Additionally, the variant is also characterized by another transition the causes an uninterrupted CCG repeat, encoding proline (i.e., CCG-CC**A** to CCG-CC**G**) occurring in complete linkage disequilibrium with the CAA to CAG transition. Notably, the last two glutamines, encoded by CA**A**-CAG in the reference HD alleles and CA**G**-CAG in LOI carriers are not included in sizing calculations in current diagnostic tests. LOI carriers in the same repeat class as reference interrupting sequence would therefore have identical polyglutamine tract lengths, but two additional uninterrupted CAG residues (Figure 1). A similar effect is observed in these individuals on uninterrupted CCG and proline repeat lengths.

**Figure 1.**
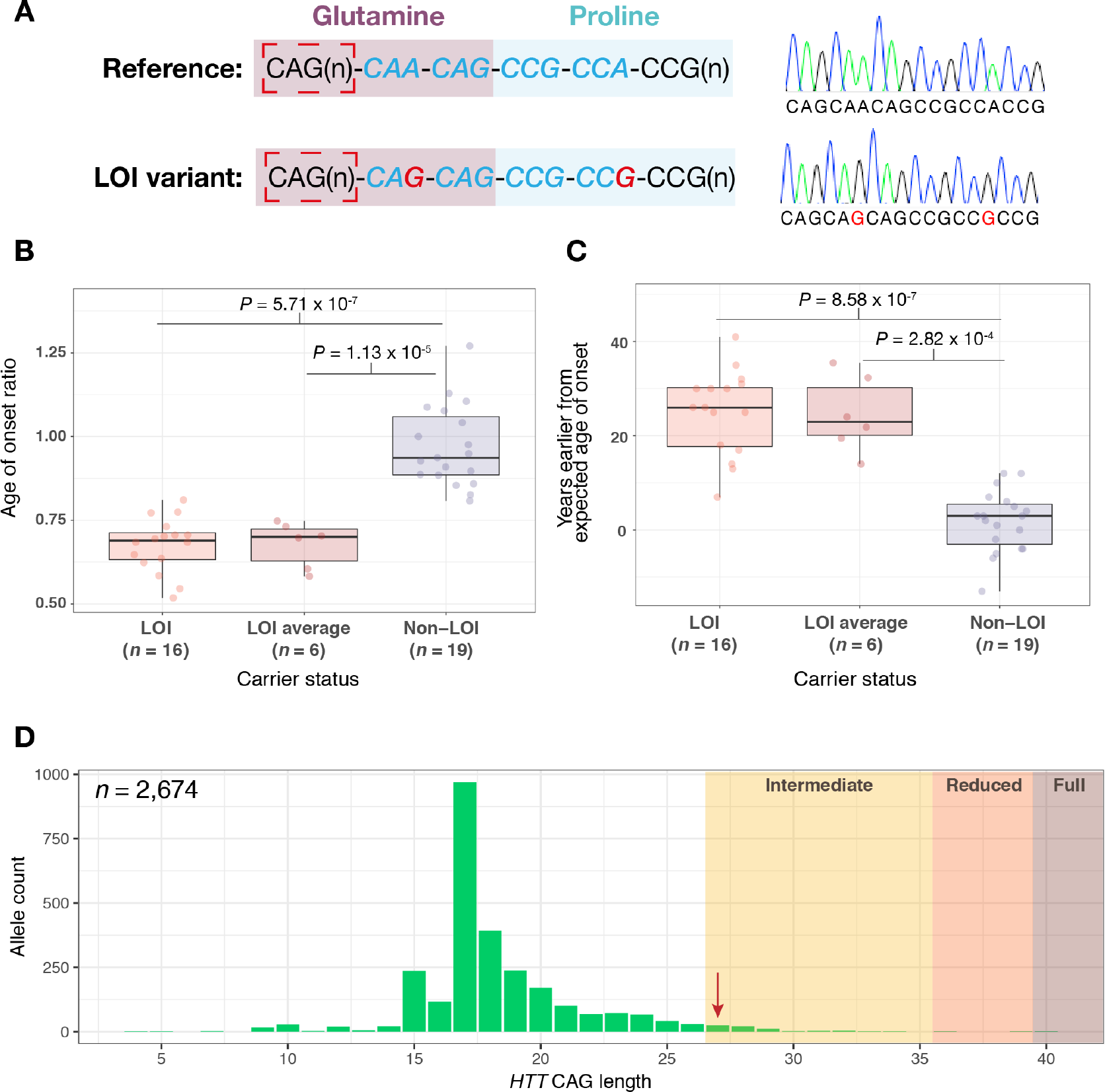
The loss of interruption (LOI) variant is associated with an earlier age of onset (AOO) in HD patients and occurs on expanded *HTT* CAG alleles. **(A)** The *HTT* CAG-CCG interrupting sequence and representative Sanger electropherograms for the reference sequence and LOI variants. The interrupting sequence is depicted in blue italic font and red nucleotides show point mutations in this region that can result in the LOI variant. The dashed red box indicates the CAG repeat that is measured in diagnostic assays for HD. Nucleotides encoding the glutamine (i.e., CAG/CAA) and proline (i.e., CCG/CCA) tracts are shaded to show that the LOI variant alters the number of contiguous CAG repeats but not the number of glutamine residues in patients. **(B)** The LOI is associated with earlier AOO as determined by the AOO ratio. **(C)** LOI carriers present with HD approximately 25 years earlier than predicted on average compared to current models for prediction of AOO. These calculations were performed using data from all HD-LOI subjects (*n*=16) as well as mean values for each HD-LOI pedigree (*n*=6), versus HD subjects with the reference interrupting sequence (*n*=19). **(D)** Distribution of the *HTT* CAG repeat lengths in the general population ascertained through genotyping from whole genome sequencing data (*n*=2,674 alleles). The LOI allele was detected in one research participant and was found on an intermediate allele (indicated with an arrow). Intermediate, reduced penetrance and fully penetrant alleles are shaded.

We identified 16 symptomatic HD subjects from six pedigrees of European ancestry from five countries (Australia, Canada, Italy, United States and the Netherlands) with this LOI variant (Figure 1, Supplementary Figure S1, mean CAG length = 39). Notably, 12 of the 16 clinically manifesting HD-LOI subjects (75%), from five of the six pedigrees, carried RP alleles (i.e., CAG 36-39). The remaining four LOI subjects were found in three of the six families and carried the LOI variant on fully penetrant HD alleles. On average, LOI carriers (*n*=16) presented with HD 25 years earlier than model predictions,^13^ which was significantly different from HD subjects with the reference interrupting sequence (i.e., CA**A**-CAG-CCG-CC**A**; *n*=19, *P*=8.58 × 10^−7^, Figure 1). Strikingly, all HD-LOI subjects presented with an extremely early AOO based on their CAG repeat length (<10th percentile of predicted AOO for CAG repeat length), and displayed a significantly lower AOO ratio compared to reference interrupting sequence subjects (*P*=5.71 × 10^−7^).

Analysis of general population controls that passed quality control via whole genome sequencing (*n*=1,337) revealed that the LOI variant is rare in unaffected individuals (minor allele frequency=0.04%, i.e., 1 in 2,674 alleles), with only one general population LOI variant detected, occurring on an intermediate CAG allele (Figure 1D). In this general population cohort, the LOI variant was therefore present in 1 of 69 intermediate alleles (IA, i.e., minor allele frequency=1.45%) and found exclusively on alleles with expanded CAG ranges (i.e., ≥CAG 27) in this study. This agrees with routine clonal sequencing of this region that has been performed by our group, where no LOI carriers have been detected in 235 unexpanded normal alleles assessed to date.

We further screened all RP individuals in the UBC HD Biobank with CAG repeat lengths in the 36-38 range (*n*=45), revealing that 60% of the clinically manifest RPs (*n*=15) versus only 7% of the asymptomatic RPs (*n*=30) carried the LOI variant in this range (*P*=2.23 × 10^−4^). Among unrelated symptomatic RP allele pedigrees in the CAG 36-38 range, 40% carried the LOI. Remarkably, when assessing RPs that presented with HD extremely early in life (i.e. <10th percentile of AOO ratio, *n*=9), 89% were LOI carriers. The LOI variant was found to occur on subsets of two common haplotypes (i.e., A1 and C1),^15^ indicating that single nucleotide variants may be unlikely to predict LOI status in HD. In contrast, previously published research^15^ has shown that the A1 haplotype is associated with elongated CAG repeat lengths.^20^

Finally, we identified a distinct variant in this region that results in a longer interrupting sequence, through the insertion of a duplicate CAA-CAG motif [i.e., 18 bp; (CAA-CAG)_2_-CCG-CCA, Supplementary Figure S2]. In the two pedigrees where this variant was present, carriers (*n*=6 HD subjects) presented 4.8 years later AOO than expected in comparison to reference interrupting sequence HD subjects (AOO percentile 60th-85^th^, AOO ratio *P*=0.04). This variant was found on a C2 *HTT* haplotype^15^ in all carriers.

### The HTT CAG-CCG LOI variant, resulting in an uninterrupted CAG tract, is associated with increased somatic and germline instability

The LOI variant was associated with increased CAG repeat tract instability (Figure 2, Table 1), both in the analysis of the somatic expansion ratio (*P*=5.39 × 10^−7^) and small-pool PCR of sperm (*P*=0.002). As expected, the somatic *HTT* CAG expansion ratio was strongly associated with CAG repeat length (*P*=4.34 × 10^−14^). In addition to LOI status and CAG repeat length (*P*=1.8 × 10^−31^), increased age was also associated with increased expansions and total instability in the small-pool sperm analyses (Table 1).

**Table 1.** Statistical analysis of somatic and germline measures of *HTT* CAG instability. The *HTT* CAG-CCG loss of interruption (LOI) was associated with increased instability in these semi-quantitative analyses. Increased donor age was also associated with increased germline instability.

| Trait | Variable | β-coefficient | P-value |
| --- | --- | --- | --- |
| Expansion frequency<br>(small-pool PCR<br>in sperm)<sup>a</sup> | CAG repeat length | 0.19 | $1.6 \times 10^{-35}$ |
|  | Age | 0.01 | 0.01 |
|  | LOI (non-LOI status) | 0.94 | 0.001 |
| Expansion ratio<br>(genomic DNA from<br>whole blood)<sup>a</sup> | CAG repeat length | 0.16 | $4.3 \times 10^{-14}$ |
|  | Age | 0.003 | 0.36 |
| | LOI (non-LOI status) | 0.57 | $5.4 \times 10^{-7}$ |

<sup>a</sup>log transformed

**Figure 2.**
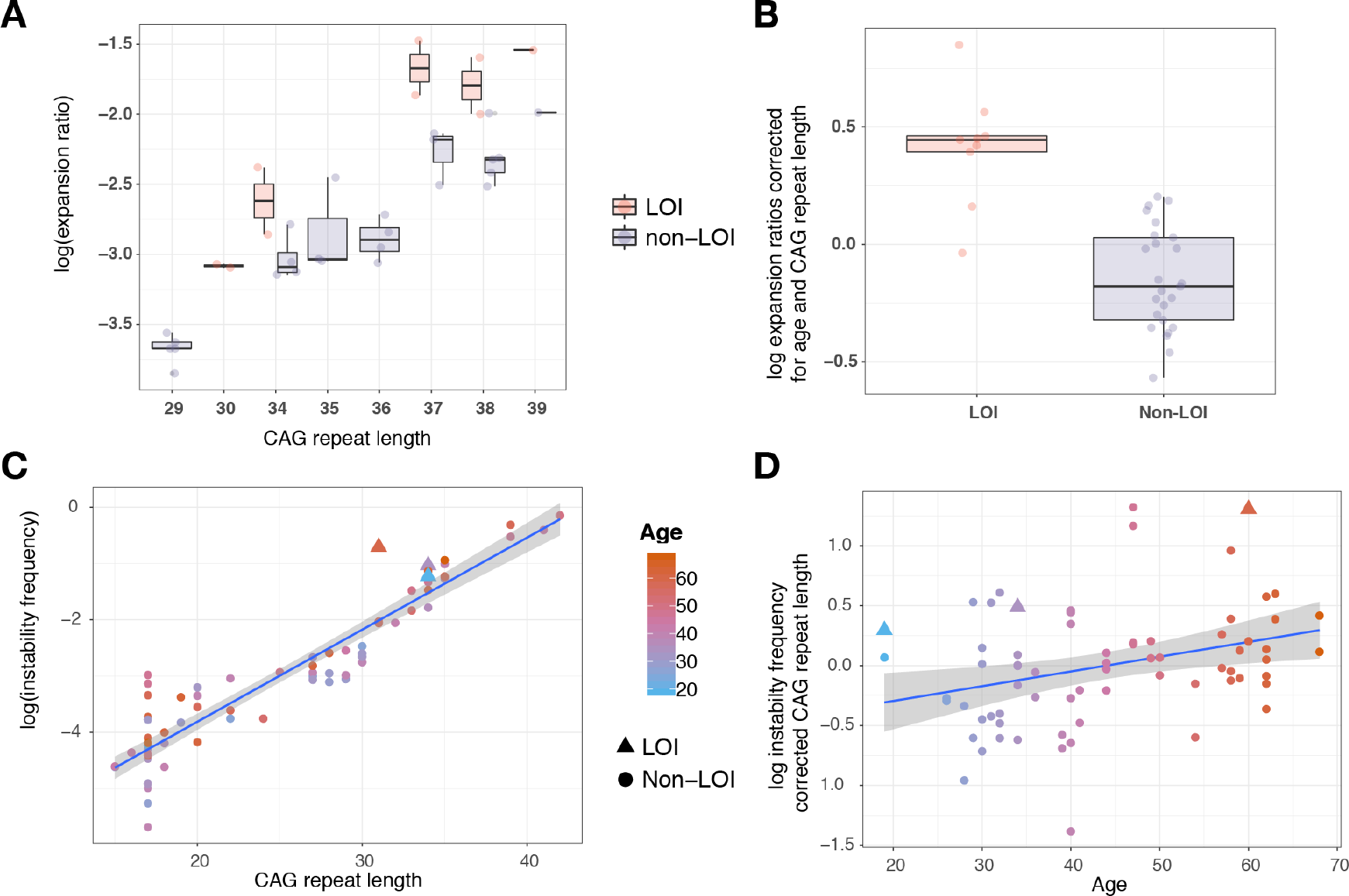
The *HTT* CAG-CCG loss of interruption (LOI) is associated with an increased frequency of CAG expansions and instability. **(A)** Expansion ratio by CAG separated by LOI carrier status. **(B)** Expansion ratio differences after correction for age and CAG repeat length between LOI and non-LOI. **(C)** Exponential relationship between somatic instability frequency and CAG repeat length, measured by small-pool PCR (variance explained by progenitor CAG repeat length, R^2^ = 0.87) **(D)** Instability frequency corrected for CAG length showing the effect of age (variance explained by age, R^2^ = 0.11). Instability for LOI subjects are indicated (one of the LOI HD patients was sampled at two separate time points). Points are colored by age at time of sampling in **(C)** and **(D)**.

## DISCUSSION

The LOI variant in the CAG-CCG interrupting sequence of *HTT* is a novel modifier of AOO of HD and is associated with increases in both somatic and germline instability of the pathogenic CAG repeat. This important finding is further supported by the fact that this variant displays familial aggregation as an autosomal dominant trait and all carriers presented with HD extremely early in life (Supplementary Figure S1). This suggests that this the LOI sequence variant is the major contributor towards differences from predicted AOO in these individuals and families. The variant is particularly relevant for HD patients in the RP range, a group of individuals that have been excluded from previous large-scale HD modifier studies.^9^ Identifying a highly-penetrant modifier variant such as the LOI variant provides an explanation for why a subset of RP patients with clinical HD manifest with the disorder much earlier than others. Here we have shown that AOO of HD is significantly influenced by the length of the uninterrupted CAG tracts and that may be more informative for AOO predictions than the polyglutamine length which would be the same for a given CAA/CAG repeat class.

Despite being rare, the LOI variant has a large effect size compared to other HD modifiers. For example, the previously-identified GWAS modifier with the largest known effect size in HD, rs146353869 in *FAN1* (2% minor allele frequency), leads to a 6.1-year alteration in AOO on average.^9^ The frequency of the LOI in the fully penetrant HD allele range still needs to be determined. However, this was indirectly assessed by a study performed in the diagnostic setting examining *HTT* null alleles, which may occur due to the inability of primers to bind to alleles carrying the LOI variant.^10^ This study described three manifest HD pedigrees that carried the LOI variant, representing 3.3% of the HD families investigated. Remarkably, one of these clinically manifest HD patients carried an IA (CAG 35) and a second pedigree was composed of RP patients with clinical HD,^10^ lending further support to the variant’s impact on AOO in RP carriers. In the past, a dilemma has been the rare clinical manifestation of HD in persons with less than 36 CAG repeats.^20^ Here, we provide a scientific basis for one such HD subject with a CAG of 35, explaining why persons in this situation could manifest with signs and symptoms of HD. The transition of penultimate CAA to CAG in this tract results in increased somatic mosaicism in blood and sperm, likely leading to increased CAG expansion in the brain and consequently earlier clinical onset than expected for identical polyglutamine tracts encoded by the canonical CAG repeat interruption sequence. This valuable information has important implications for the diagnosis of HD in similar patients.

In this study, all patients carrying the LOI variant presented with HD in the earliest percentile for the corresponding polyglutamine repeat typically encoded by the reference CAG repeat and interruption. The variant was predominantly seen in RP patients with clinical HD, making up 75% of clinically symptomatic carriers. However, a strong effect was still observed in HD patients with fully penetrant alleles, indicating that the finding is generalizable to the whole HD population. In our large cohort of population controls (*n*=3,314 alleles), individuals with unexpanded alleles did not carry the LOI (*n*=3,242 alleles, i.e., <27 CAG), indicating that the variant is more prevalent at longer CAG repeat lengths, possibly due to a higher likelihood of CAG expansion.^21-23^

In addition to the LOI variant, we found an interrupting sequence polymorphism that is associated with clinically meaningful later AOO and is characterized by an extra CAA-CAG motif at the end of the glutamine tract (Supplementary Figure S1). This lends further support to the role of somatic instability in HD as the variant may increase repeat stability by preventing slippage during DNA replication. The DNA sequence changes that cause the LOI variant result in a longer pure CAG repeat length; however, at the protein level, they do not alter the number of pathogenic glutamine amino acids, since CAA and CAG both encode glutamine residues (Figure 1). This indicates that the length of the uninterrupted CAG tract may be a more informative measure for use in predictive models than polyglutamine length.

The modifying effect on AOO must therefore occur through a mechanism upstream from translation of the mutant protein. Our analyses indicate that somatic instability, resulting in a mosaic of longer CAG repeat and polyglutamine tracts in vivo, is the likely mechanism for modification of HD onset by the LOI variant as shown by two orthogonal methods. Future predictive models of AOO should therefore consider assessing the number of uninterrupted CAG residues, rather than effective polyglutamine lengths, in their estimates.

This is particularly important for those HD patients with diagnostic CAG repeat lengths in the RP range (CAG 36-39). Only a small, but significant, proportion of these individuals will go on to develop HD.^3; 25^ These findings therefore challenge the prior beliefs that polyglutamine length determines AOO and clearly demonstrate that length of the uninterrupted CAG length, and not polyglutamine, is the major contributor to AOO in HD. Further, increased instability of the CAG repeat in HD subjects with the LOI highlights somatic mosaicism as a key contributor to the pathogenesis of HD.

Modification of clinical presentation by loss of glutamine-encoding CAA interruptions to pure CAG repeats has been reported in other polyglutamine disorders. For example, loss of interrupting CAA codons within the polyglutamine-encoding repeats of *ATXN2* and *TBP* have been shown to modify the pathogenicity and onset of two spinocerebellar ataxias (SCAs), SCA2 and SCA17.^25-28^ Our finding that loss of the reference CAA interruption hastens age of onset in HD, and that an extra CAA interruption conversely delays onset, expands the number of polyglutamine disorders where variable CAG and CAA repeat composition can result in phenotypic differences without alteration of translated polyglutamine length. Further, our study suggests that rare variations in polyglutamine codon structure may be present in patient populations of other polyglutamine diseases and could account for phenotypic outliers in those conditions. Future studies should investigate other polyglutamine repeat disorders by recruiting subjects with known AOO or other phenotypic characteristics and sequencing the repeat tracts directly to determine trinucleotide composition.

Somatic instability in HD has returned to the fore with recent HD onset GWAS uncovering the importance of DNA repair genes.^8; 9^ Prior to these genomic studies, previous work by our group^29^ has shown that the composition of CAA interruptions in the CAG repeat may be responsible for phenotypic differences between HD mouse models, and could be similarly mediated by differences in somatic instability. Other research has shown that somatic CAG expansion rates differ across tissues in all HD patients; with the striatum being the most vulnerable to this phenomenon,^30^ and some HD patients exhibiting over 1,000 CAG repeats in this brain.^23; 31^ Somatic instability observed in blood is less pronounced and ranges within a few CAG repeat sizes of the progenitor CAG.^23^

We also demonstrate that increased age is associated with a higher frequency of CAG repeat expansions in sperm (Table 1), which has potential clinical implications in relation to the rate of new mutations from older IA fathers. We have previously shown that CAG repeat length correlates with measures of instability in sperm,^19; 32^ but the finding with regards to the potential influence of donor age on stability has not been reported. In our somatic and germline assays, the effect of the LOI variant on stability measures was larger than that of CAG repeat length, indicating the dramatic influence of this genotype on *HTT* CAG repeat instability in humans.

The LOI variant remains laborious to genotype with conventional clonal sequencing methods, making it difficult to infer the frequency of the variant in the broader HD patient populations. Further study will be necessary to determine the impact of the LOI across the fully penetrant HD CAG repeat range. Since common *HTT* haplotypes do not predict LOI status with sufficient resolution for diagnostic purposes, high throughput sequencing of the entire *HTT* gene locus in LOI carriers could aid in identifying rarer, more easily genotyped, tag variants. The feasibility of genome-wide arrays to capture such proxies should also be explored since carriers of this particular variant may not be detected in genome wide association studies with sufficient resolution. This could occur if the LOI variant has a *de novo* origin in a large proportion of carrier families. Further, the LOI variant may result in diagnostic errors through drop out in current diagnostic sizing of the CCG repeat.^10^ Our analysis of whole genome sequencing information indicates that genotyping the interrupting sequence is feasible and that more targeted approaches, using similar technologies, should be explored for clinical genetic testing applications.

In conclusion, we have described a novel modifier of HD clinical onset that has a larger impact than all previously identified modifier variants, most dramatically observed in the RP range. This provides conclusive support for the role of somatic repeat instability and DNA repair in modifying HD AOO. The relevance of somatic mosaicism in HD was first documented 15 years ago,^22; 23^ with the greatest instability observed in the brain and particularly those regions most pertinent to HD pathogenesis. Our study therefore provides an explanation as to why a proportion of RP carriers present with HD signs and symptoms early in their lifetime. This may have substantial implications for clinical practice and could provide important information for individuals that present with RP alleles. This LOI is present at high frequency in symptomatic RPs and may therefore provide additional information for genetic counselling of subjects with CAG repeats at the lower end of the HD range.

## ACKNOWLEDGEMENTS

This work was supported by a Canadian Institutes of Health Research Foundation Grant awarded to M.R.H. S.W.S. is the GlaxoSmithKline-CIHR Chair in Genome Sciences. We would like to thank the Centre for Molecular Medicine Therapeutics and BC Children’s Hospital Research Institute, as well as The Centre for Applied Genomics at the Hospital for Sick Children and the University of Toronto McLaughlin Centre for support. Additionally, we would like to acknowledge Léal Makaroff for his assistance with developing and validating assays for somatic instability. We also wish to thank all the HD patients and families worldwide who have chosen to participate in research, including those from the Centre for Huntington Disease and the HD BioBank at UBC; Leiden University Medical Center, the Netherlands; the Children’s Hospital in Westmead, Sydney, Australia; as well those collected through with the support of Lega Italiana Ricerca Huntington (LIRH) Foundation in Rome, Italy. Without the support of HD families none of this research would be possible.

