## Supplementary Figures S1 and S2 for "Length of uninterrupted CAG repeats, independent of polyglutamine size, results in increased somatic instability and hastened age of onset in Huntington disease"

[1]Centre for Molecular Medicine Therapeutics, Department of Medical Genetics, University of British Columbia, Vancouver, BC, Canada; [2]Illumina Inc, San Diego, California, USA; [3]Department of Clinical Neuroscience, Karolinska Institutet, Stockholm, Sweden; [4]Department of Psychiatry, University of British Columbia, Vancouver, BC, Canada; [5]The Hospital For Sick Children, The Centre for Applied Genomics, Genetics and Genome Biology; [6]University of Toronto, Department of Molecular Genetics; [7]Department of Clinical Genetics, Children's Hospital at Westmead; [8]Department of Clinical Genetics, Leiden University Medical Center, Leiden, the Netherlands; [9]Huntington and Rare Diseases Unit, Fondazione IRCCS Casa Sollievo della Sofferenza, San Giovanni Rotondo, Italy; [10]McLaughlin Centre, University of Toronto.

**Supplementary Figure S1 HD loss of interruption (LOI) patient pedigrees included in the current study (*n*=6).** Affection status, CAG sizes, age of onset information and LOI genotype are indicated

**HD-LOI-01**

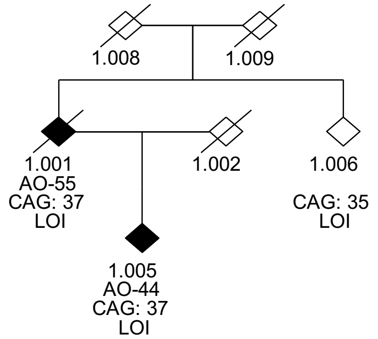


**HD-LOI-02**

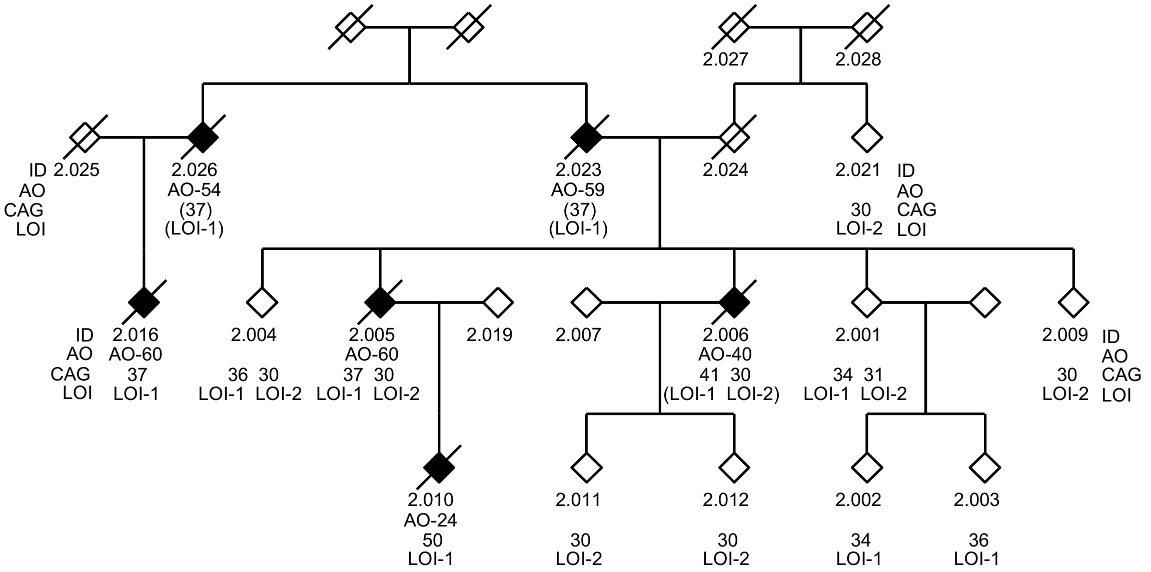

**HD-LOI-03**

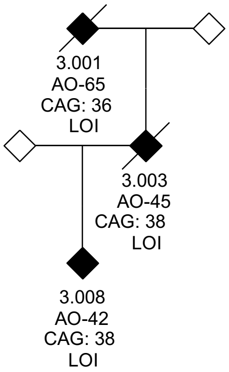


**HD-LOI-04**

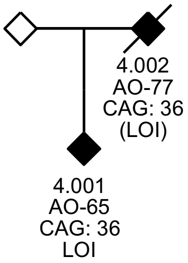


**HD-LOI-05**

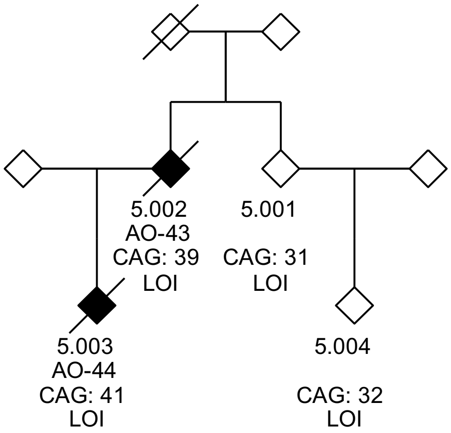

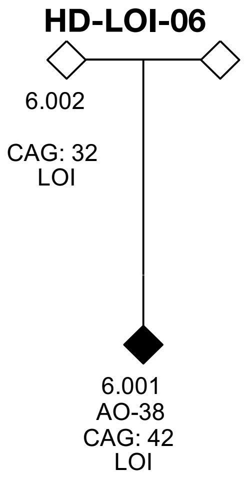
HD-LOI-06
6.002
CAG: 32
LOI
6.001
AO-38
CAG: 42
LOI

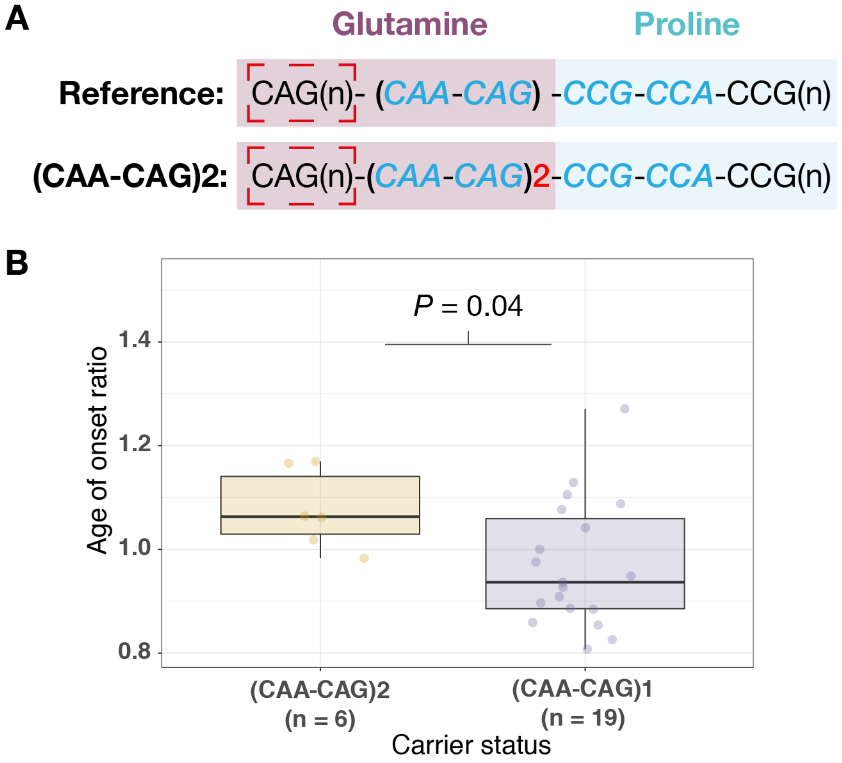


**Supplementary Figure S2. An interrupting sequence variant that results in an additional CAA-CAG repeat is associated with a later age of onset in HD patients. (A)** The reference *HTT* CAG-CCG interrupting sequence in relation to the $(CAA\text{-}CAG)_2$ variant. The dashed red box indicates the CAG repeat that is measured in diagnostic assays for HD. Nucleotides encoding the glutamine (i.e., CAG/CAA) and proline (i.e., CCG/CCA) tracts are shaded. **(B)** The $(CAA\text{-}CAG)_2$ variant is associated with later age of onset (AOO) as determined by the AOO ratio in HD subjects compared to HD subjects with the reference interrupting sequence ($n$=19). $(CAA\text{-}CAG)_2$ carriers present on average 4.8 years later than the majority of HD patients with the reference CAG repeat interruption.
